# Molecular evolutionary consequences of island colonisation

**DOI:** 10.1101/014811

**Authors:** Jennifer E. James, Robert Lanfear, Adam Eyre-Walker

## Abstract

Island endemics are likely to experience population bottlenecks; they also have restricted ranges. Therefore we expect island species to have small effective population sizes (*N_e_*) and reduced genetic diversity compared to their mainland counterparts. As a consequence, island species may have inefficient selection and reduced adaptive potential. We used both polymorphisms and substitutions to address these predictions, improving on the approach of recent studies that only used substitution data. This allowed us to directly test the assumption that island species have small values of *N_e_*. We found that island species had significantly less genetic diversity than mainland species; however, this pattern could be attributed to a subset of island species that had undergone a recent population bottleneck. When these species were excluded from the analysis, island and mainland species had similar levels of genetic diversity, despite island species occupying considerably smaller areas than their mainland counterparts. We also found no overall difference between island and mainland species in terms of effectiveness of selection or mutation rate. Our evidence suggests that island colonisation has no lasting impact on molecular evolution. This surprising result highlights gaps in our knowledge of the relationship between census and effective population size.

## Introduction

Island species have long been considered to be under greater threat of extinction than their mainland counterparts (Mckinney 1997; Frankham 1997; Johnson & Stattersfield 1990; Jones et al. 2003; Purvis et al. 2000). Although extinction itself is caused by a number of stochastic factors, not least human activity (Burgess et al. 2013; Pimm et al. 1988), the susceptibility of island populations may also be a consequence of population genetics. Island species are likely to have experienced population bottlenecks at some point in their evolutionary history due to founder events during the initial island colonisation. As only a fraction of individuals from the original population found an island population, only a fraction of the original genetic diversity of the population will be maintained, and effective population sizes (*N_e_*) will be small (Nei et al. 1975). In addition, island species are restricted to relatively small areas, which could impose long-term restrictions on census population sizes, and in turn on long-term *N_e_*. Therefore it may be that island species are genetically vulnerable.

Low diversity/low *N_e_* could theoretically reduce the adaptive potential of a species, as standing levels of genetic variation determine the alleles that are immediately available for evolution to act upon (Messer & Petrov 2013; Barrett & Schluter 2007; Hermisson & Pennings 2005). In addition, populations founded by a small number of individuals will experience increased inbreeding. Inbreeding results in an increasingly homozygous population, and therefore there is a greater risk that deleterious recessive alleles will be exposed (Charlesworth & Charlesworth 1987), which could have significant fitness costs. There is some evidence that bottlenecked species do experience a loss of fitness: for example, Frankham et al. (Frankham et al., 1999) demonstrated that laboratory populations of *Drosophila* showed reduced evolvability (in terms of ability to tolerate increasing concentrations of an environmental pollutant) after a bottleneck; while Briskie and Mackintosh (Briskie & Mackintosh 2003) uncovered a link between the severity of population bottlenecks and loss of fitness in birds.

In addition, species with low effective population sizes are expected to have inefficient selection, resulting in high levels of deleterious mutations segregating and a tendency to fix deleterious mutations. However, past studies investigating the differences in the efficiency of selection between island and mainland species have provided only limited support for this prediction. Johnson and Seger (Johnson & Seger 2001) found some evidence that island species had less efficient selection, but this was for a small and taxonomically restricted dataset. Woolfit and Bromham (Woolfit & Bromham 2005) used a much larger and more varied dataset, however, they reported a difference between island and mainland species that was only significant at the one-tailed level, while Wright et al. (Wright et al. 2009) found no significant difference between island and mainland species. This may be because previous studies have considered rates of substitution, usually the ratio of the rate of non-synonymous substitution to the rate of synonymous substitution (ω). The problem with considering substitution data is that a reduction in *N_e_* is expected to increase the rate at which slightly deleterious mutations are fixed, but reduce the rate at which advantageous mutations are fixed, particularly if the rate of adaptation is limited by the supply of mutations. We therefore cannot make a clear prediction about the effect of *N_e_* on substitutions. This issue can be addressed by using polymorphism data instead of substitution data, using the number of nonsynonymous polymorphisms divided by synonymous polymorphisms, because advantageous mutations are not expected to significantly contribute to polymorphism (Kimura 1984; Kryazhimskiy & Plotkin 2008; Ho et al. 2011).

It seems likely that adaptive evolution might occur for at least some island species, despite their predicted low effective population sizes, due to the fact that the species is encountering a novel habitat. Although populations with large effective population sizes may have more efficient selection, we might also expect positive selection to play a significant role after a colonisation event, as species adapt to new environmental requirements and ecological niches. However, in making predictions regarding adaptive evolution it is important to consider the direction of colonisation. Although island species most commonly colonise an island from a nearby mainland, occasionally lineages that originated on islands re-colonise a mainland, providing an interesting contrast in terms of molecular evolution. Species colonising mainlands from islands are likely to experience population size increases, and therefore increases in *N_e_*. This could result in a spate of rapid molecular evolution in the new mainland population as advantageous mutations that were previously effectively neutral become fixed (Takano-Shimizu 1999; Charlesworth & Eyre-Walker 2007).

However, predictions about the molecular evolution of island species are predicated on the crucial assumption that island species do in fact have lower *N_e_* and levels of genetic diversity than mainland species. Whether this is in fact the case is not certain, because census population size can sometimes be a poor indicator of genetic diversity (Lewontin 1974; Bazin et al. 2006; Leffler et al. 2012; Romiguier et al. 2014). Although some studies uncover a link between the two (for overview, see (Frankham 2012)), other authors have not found a relationship; for example, Nabholz et al. (Nabholz, Mauffrey, et al. 2008) did not find a strong relationship between mammalian mitochondrial diversity and life-history traits associated with *N_e_* (such as body mass), or between diversity and IUCN category, an index partly based on assessments of census population size. More generally, there is surprisingly little variation in levels of diversity between species; one recent paper reported a range of nucleotide diversities of only 800-fold across a range of taxa, many orders of magnitude smaller than their estimated census population size differences (Leffler et al. 2012). The determinants of genetic diversity remain poorly understood.

One possible complicating factor is the mutation rate. Both Nabholz et al. (Nabholz, Glémin, et al. 2008) and Romiguier et al. (Romiguier et al. 2014) found evidence suggesting that there are lineage-specific differences in the mutation rate, in mitochondrial and nuclear data respectively. How the mutation rate evolves is contentious: if selection is responsible for determining the mutation rate, populations with high effective population sizes should have the lowest mutation rates, because selection will be more effective at reducing the rate: this is because whether a mutation can be selected depends on the strength of selection being over 1/*N_e_*. However, support for this prediction remains mixed. For example, in previous studies of island-mainland systems (all of which controlled for phylogenetic non-independence), two found no difference in substitution rate between island and mainland lineages (Woolfit & Bromham 2005; Johnson & Seger 2001), while another found that it was mainland species that had higher rates of substitution (Wright et al. 2009), the opposite of what we might expect if the mutation rate depends on the population size. Another factor that may contribute to unexpected patterns of diversity is selection at linked sites: this reduces genetic diversity, particularly in genomic regions with low rates of recombination (Frankham 2012; Gillespie 2000; Maynard Smith & Haigh 1974). Linked selection may occur more frequently in populations with high values of *N_e_*, reducing diversity more rapidly than in populations with a low *N_e_*. On the other hand, it could be that selective sweeps occur more commonly in species adapting to a new environment e.g. (Montgomery et al. 2010).

In summary, we expect island species to have low effective population sizes and because of this we expect them to have low genetic diversities. We also expect selection to be less efficient in island species, leading to higher ratios of nonsynonymous to synonymous polymorphism, and potentially to increases in the mutation rate (the mutation rate might increase to such an extent that island and mainland species have similar diversities). Whether we expect island species to have higher ratios of nonsynonymous to synonymous substitution depends on how much adaptive evolution there is, and how this is affected by effective population size and the act of colonisation. If there is no adaptive evolution then island species are expected to have higher values for ω; however, adaptive evolution could potentially be either reduced in island species because of their low *N_e_* or increased because of adaptation to a new environment, given that in most cases the island is the new environment that is colonised. Here we perform the first analysis of polymorphism data from a dataset of phylogenetically independent pairs of island and mainland species, and combine this with substitution data. The paired study design is crucial: there are a large number of life history traits that are known to influence molecular evolution (e.g. body size, fecundity, generation times) and could therefore act as confounding factors (Bromham 2011; Lanfear et al. 2013). Closely related island and mainland species have similar life-history traits, and even if there is variation it is not expected to be systematic, and so should not bias our results. Therefore, island colonisation itself will be the primary reason for any differences in molecular evolution between island and mainland species (Johnson & Seger 2001; Woolfit & Bromham 2005).

## Results

### Dataset overview

To investigate the consequences of island colonisation on molecular evolution we compiled data for 120 island-mainland comparisons. Some comparisons comprised a single island and single mainland species, while some consisted of multiple island and multiple mainland species. In the majority of cases, the inferred direction of colonisation is from mainland-to-island. The data is dominated by mitochondrial sequences from birds (Table 1a) but we have a reasonable number of mitochondrial sequence comparisons available for invertebrates and reptiles, and a moderate number of nuclear sequence comparisons. For 70 of our comparisons, multiple sequences from the same species were available, allowing us to conduct polymorphism analyses. Again, this dataset is dominated by mitochondrial data from birds (Table 1b). For a full list of species used in this analysis, and for complete details of results, please see the archived data at: http://dx.doi.org/10.6084/m9.figshare.1296151.

**Table 1a and b.**
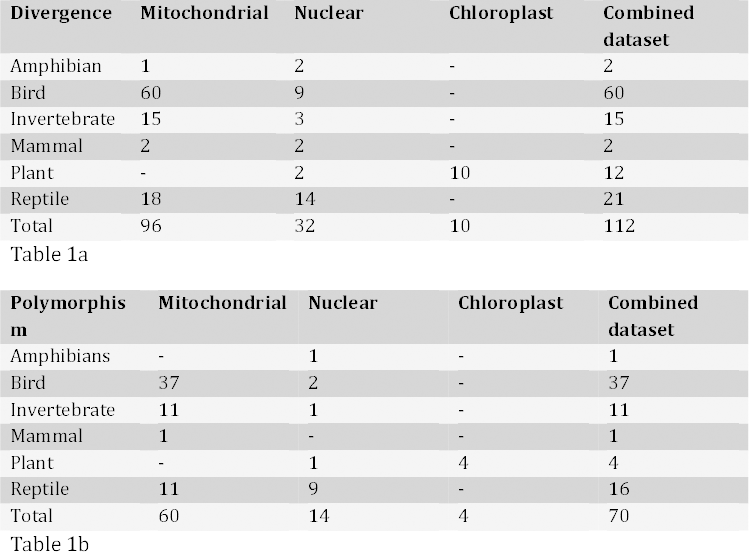
An overview of the sequences gathered in this analysis, split by DNA type and taxonomic group. For analyses that combined data across DNA types, each species comparison appeared only once: the numbers of sequences available in these cases are given in the ‘combined dataset’ column. When choosing between sequences from different genomes for a particular comparison, we always used the longest sequence.

### Geography

Island species are studied from a molecular evolutionary perspective because they are expected to have smaller populations than mainland species due to their small ranges. However, this assumption is rarely tested. In this study, the ranges of the species used were confirmed where possible using the IUCN database (IUCN 2014). The mean range of island species was 5,780 km^2^, while for mainland species this mean range was over 4,080,000 km^2^. The ratio of island to mainland range sizes did not exceed 0.25 for any of the comparisons used, and in the majority of cases island species had ranges which were less than 1% of the area of those of their mainland counterparts (Figure 1). Therefore we have evidence that the island species used in this study inhabit substantially smaller geographic regions than their mainland relatives, although we have no information on population density.

**Fig. 1.**
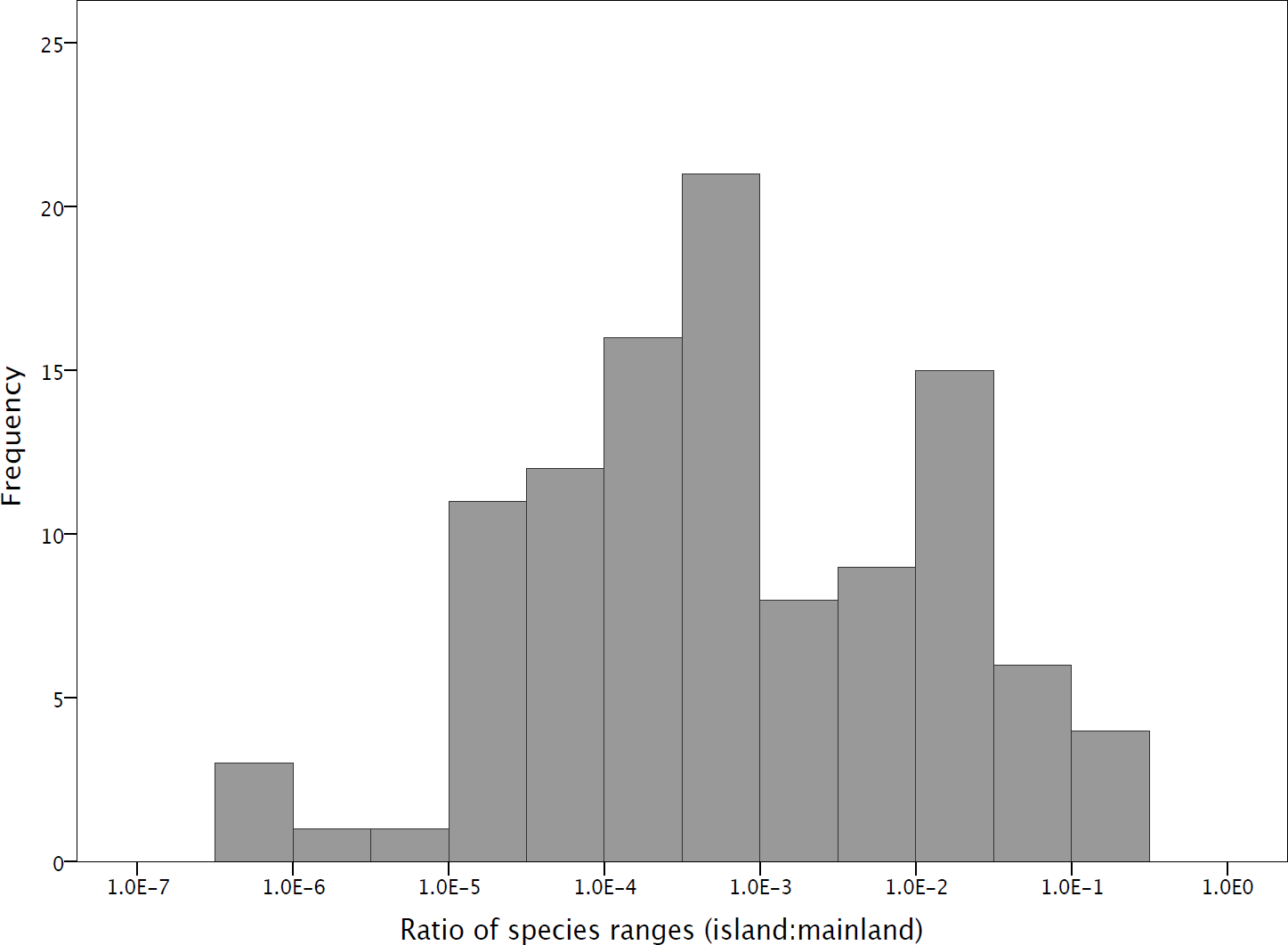
The frequency distribution of the ratios of island:mainland species range areas

### Synonymous diversity

We might expect island species to have lower diversity than their mainland counterparts for two reasons. Firstly, island species inhabit substantially smaller areas than their mainland relatives; resulting in a smaller census population size and hence potentially a smaller long-term *N_e_*. Secondly, island populations are likely to be founded by few individuals, which again is expected to result in a small *N_e_*. As expected, we find that island species have significantly lower synonymous site nucleotide diversity overall, and when we consider mitochondrial and nuclear DNA separately (Table 2). Chloroplast sequences show the opposite pattern, but as there are only 4 comparisons this is likely to be due to sampling error. When different taxonomic groups were considered separately, island birds had significantly lower levels of diversity than mainland birds, while for both reptiles and invertebrates there was no significant pattern (Table 2) (for other groups we do not have enough data to make a valid comparison). However, despite being statistically significant, the differences between mainland and island species are relatively modest. Mainland species have on average 40% more diversity than island species, and in about one third of cases, island species have higher diversity than their mainland relatives.

**Table 2.** Differences in synonymous nucleotide diversities (π_s_) between island and mainland species. The number of comparisons used in each analysis is given in the second column (n), with the significance level of the Wilcoxon signed-ranks test given in the last column. Each particular species comparison appears only once in each dataset.

| Dataset | n | Mean Island $\pi_s$ | Mean Mainland $\pi_s$ | Larger ranks-Island:Mainland | p-value |
| --- | --- | --- | --- | --- | --- |
| Combined | 70 | 0.0270 | 0.0388 | 19:43 | 0.010 |
| Chloroplast | 4 | 0.00231 | 0.000575 | 2:0 | 0.180 |
| Mitochondrial | 60 | 0.0320 | 0.0524 | 19:39 | 0.014 |
| Nuclear | 14 | 0.00147 | 0.00686 | 1:7 | 0.036 |
| Bird mitochondrial | 37 | 0.0117 | 0.0283 | 11:25 | 0.012 |
| Invertebrate mitochondrial | 11 | 0.0784 | 0.0582 | 4:6 | 0.646 |
| Reptile mitochondrial | 11 | 0.0562 | 0.116 | 4:7 | 0.131 |

It is potentially possible to differentiate between the two possible causes of the lower diversity in island species by considering the ratio of island to mainland nucleotide diversity as a function of the divergence between the island and mainland species. In this analysis we use the total number of island and mainland synonymous substitutions (dS) as an estimator of species divergence, however it should be noted that this is a crude estimator as dS is dependant on both generation time and mutation rate. If most of the reduction in diversity is due to a bottleneck during colonisation, then we expect the ratio of island to mainland diversity to be greatest when the evolutionary divergence is longest. In contrast, if diversity is largely determined by population sizes after colonisation then we might expect the ratio of island to mainland diversity to decline with evolutionary divergence. Consistent with the bottleneck hypothesis, we find that π_s_(island)/(π_s_(island)+π_s_(mainland)), the normalised level of neutral island diversity, is positively correlated to the total number of synonymous substitutions between island and mainland species (Pearsons correlation r= 0.318, p= 0.012) (Figure 2). The correlation increases in strength if we restrict the analysis to mainland-to-island colonisation events (r = 0.384, p=0.004), and is negative, though non-significant, if we consider colonisations that occurred in the opposite direction (r = −0.129, p = 0.74). This positive correlation appears to be driven by a group of island species/clades that are recent colonists and have no synonymous diversity (Figure 2), because the positive correlation disappears when these species are removed (r = 0.214, p = 0.150). Although the low levels of diversity we have recorded could be a result of low levels of mutation and/or short sequences, this explanation is unlikely because we would expect equal numbers of island and mainland species to have low diversity (i.e. in Fig 2 we would expect an equal number of points clustering at 1 on the y-axis as at 0), which is not what we observe.

**Fig 2.**
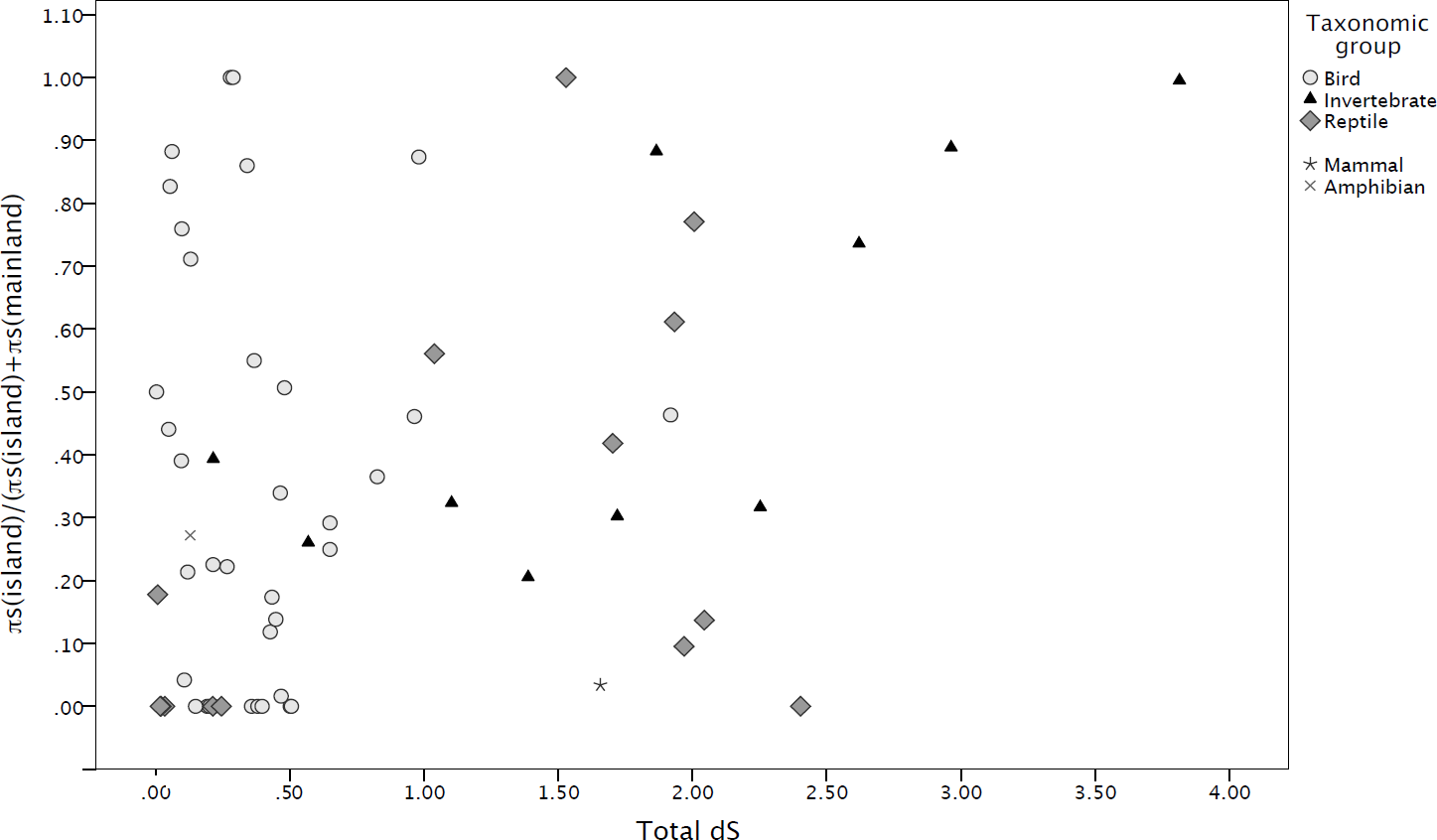
The ratio of island diversity to the combined island and mainland diversity, π_s_(island)/( π_s_(island)+ π_s_(mainland), where π_s_ is synonymous diversity, plotted against total divergence (dS) between island and mainland species.

Reptiles are disproportionately represented amongst the species with no genetic diversity in the island species/clades (6 out of 14 reptiles compared to 9 out of 36 birds and 0 out 9 invertebrates). If each phylogenetic group is considered individually we find a significant positive correlation between π_s_(island)/(π_s_(island)+π_s_(mainland)) and dS for invertebrates (r = 0.752, p = 0.012) and positive but non-significant correlations for birds and reptiles (figure 2) (we do not have enough data to study the other groups individually). As a group, birds appear to retain the highest levels of diversity, with some species seemingly not undergoing a population bottleneck during the colonisation event, perhaps because there are more individuals initially founding the island population and/or because there is continued migration from the mainland. This is compatible with the greater dispersal ability of birds compared to other animal groups. Reptiles on the other hand appear to experience a quite severe loss of diversity during founder events.

Although our results are consistent with the idea that the genetic diversity of island species is able to recover over time, either through continued immigration or the accumulation of new genetic diversity *in situ*, an alternative interpretation is that island species that are not diverse simply go extinct. This may be why only young species have low levels of diversity (out of 62 comparisons, only the chameleon *Archaius tigris* was moderately divergent without any synonymous diversity at all). These explanations are not necessarily mutually exclusive. Nevertheless it is surprising that aside from those species with no synonymous diversity, in most cases island species have similar and in some cases more genetic diversity than their mainland counterparts. If we remove the comparisons in which island diversity is zero and re-analyse the data we find that the remaining island species do not have lower synonymous diversity than mainland species (Wilcoxon signed-ranks test, n = 48, p = 0.258). This suggests that island species/clades only have lower levels of diversity if they have recently (in terms of generations) undergone a population bottleneck.

### Effective population sizes

The fact that the genetic diversity of island species is generally not lower than that of mainland species suggests that they do not have lower effective population sizes. To investigate this, we estimated effective population size by dividing synonymous diversity by synonymous divergence (using synonymous divergence to approximate mutation rate) and compared island species to their mainland counterparts. Note that these effective population size estimates can only be compared against each other, since we are dividing the diversity by the product of the mutation rate per generation and the number of generations since the mainland and island species diverged. Mainland species had significantly greater effective population sizes than island species overall (Wilcoxon signed-ranks test, n = 66, p = 0.030); however, if we exclude those comparisons in which the island species had no synonymous diversity, the difference between island species and mainland species is no longer significant (n = 45, p = 0.281).

### Efficiency of selection

Selection is expected to be less efficient in species with small *N_e_*. However, we have found little evidence to suggest that island species have lower long-term effective population sizes than mainland species. It is therefore perhaps not surprising that we find little evidence for selection being less efficient in island species. Using polymorphism data we compared π_n_/(π_n_+π_s_) between island and mainland species and found no significant differences between island and mainland species/clades (Wilcoxon signed-ranks test, n = 51, p = 0.389); we also found no difference when considering different DNA types separately, although when splitting by taxonomic group the difference between island and mainland bird species is just significant (Table 3). It should be noted however that most of the island species that have no synonymous polymorphisms also have no non-synonymous polymorphisms and hence are excluded from the analysis because π_n_/(π_n_+π_s_) is undefined.

**Table 3.** Differences in π_n_/(π_n_+π_s_) between island and mainland species. The number of comparisons used in each analysis is given in the second column (n), with the significance level of the Wilcoxon signed-ranks test given in the last column. Each particular species comparison appears only once in each dataset.

| Dataset | n | Mean Island $\pi_n/(\pi_n+\pi_s)$ | Mean Mainland $\pi_n/(\pi_n+\pi_s)$ | Larger ranks-Island:Mainland | p-value |
| --- | --- | --- | --- | --- | --- |
| Combined | 51 | 0.175 | 0.093 | 25:23 | 0.389 |
| Chloroplast | 1 | 0.257 | 0.223 | 1:0 | - |
| Mitochondrial | 48 | 0.170 | 0.092 | 23:22 | 0.569 |
| Nuclear | 3 | 0.182 | 0.126 | 1:2 | - |
| Bird | 30 | 0.268 | 0.103 | 17:11 | 0.050 |
| Invertebrate | 10 | 0.035 | 0.055 | 5:5 | 0.646 |
| Reptile | 20 | 0.027 | 0.095 | 2:5 | 0.063 |

We also find no significant differences between island and mainland species for ω (nonsynonymous divided by synonymous divergence) overall, or if we split the data by phylogenetic group or genome type (Table 4). However, there is an expectation that ω will increase during a population size expansion (Charlesworth & Eyre-Walker 2007) and so we might expect island-to-mainland colonisations to show different patterns to mainland-to-island colonisations. If we restrict our analysis to mainland-to-island colonisations we still do not observe a significant difference between island and mainland ω overall, or for each genome, although if we split by phylogenetic group the result for birds is significant (Table 4). We also do not observe any significant difference in ω(mainland) / ω(island) between species that have colonised the island from the mainland, and the mainland from the island (independent samples t-test, p = 0.315), contrary to the results of Charlesworth and Eyre-Walker (Charlesworth & Eyre-Walker 2007).

**Table 4a and 4b.**
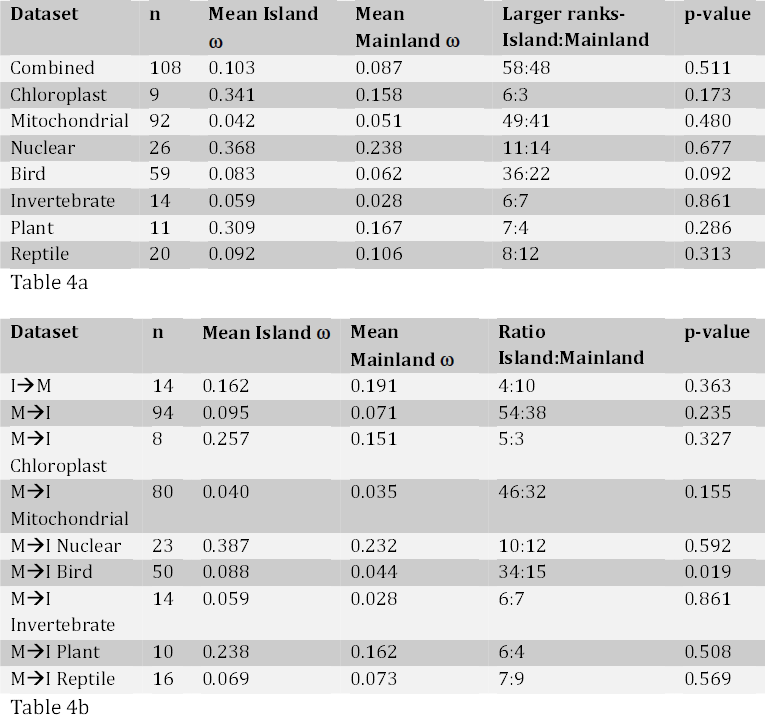
Differences in ω, (nonsynonymous divided by synonymous divergence) between island and mainland comparisons. The number of comparisons used in each analysis is given in the second column (n), with the significance level of the Wilcoxon signed-ranks test given in the last column. Each particular species comparison appears only once in each dataset. In a), the total dataset is analysed and then divided by DNA type and taxonomic group, while in b), the comparisons are split by colonisation direction; I➔M refers to comparisons in which the colonisation direction was island-to-mainland, while M➔I is mainland-to-island. Where the colonisation direction was mainland-to-island, comparisons were further divided by genome and taxonomic group

### Adaptive evolution

Colonisation of an island might be expected to lead to a burst of adaptive evolution, since the colonisers are experiencing a new environment that might have empty niches into which the species can adaptively evolve. To investigate whether colonisation leads to higher rates of adaptive evolution we estimated the rate of adaptive amino acid substitution along the island and mainland lineages using two approaches. First we calculated the direction of selection (DoS) statistic for each lineage. We find that on average the DoS is negative in both island and mainland species (Table 5), indicating that slightly deleterious mutations are prevalent in our data. We find no significant difference in values of DoS between island and mainland species, either when considering the dataset as a whole, or when the results are analysed separately depending on the direction of colonisation. However, DoS is sensitive to slightly deleterious mutations segregating in the population, and therefore any changes in the relative frequencies of deleterious mutations between island and mainland species will influence DoS, potentially masking a signal of adaptive evolution (Nielson 2005). Unfortunately, we did not have sufficient polymorphism data to correct for slightly deleterious mutations by removing low frequency polymorphisms (Fay et al. 2001; Charlesworth & Eyre-Walker 2008).

**Table 5.** Differences in DoS between island and mainland species, for the combined dataset, and for the dataset split by the direction of colonisation. The number of comparisons used in each analysis is given in the second column (n), with the significance level of the Wilcoxon signed-ranks test given in the last column. I➔M refers to comparisons in which the colonisation direction was island-to-mainland, while M➔I is mainland-to-island Differences in synonymous divergence (dS) between island and mainland species. The number of comparisons used in each analysis is given in the second column (n), with the significance level of the Wilcoxon signed-ranks test given in the last column. Each particular species comparison appears only once in each dataset.

| Dataset | n | Mean Island DoS | Mean Mainland DoS | Larger ranks: Island:Mainland | p-value |
| --- | --- | --- | --- | --- | --- |
| Combined | 50 | -0.180 | -0.167 | 25:25 | 0.783 |
| I → M | 8 | -0.144 | -0.109 | 3:5 | 0.674 |
| M → I | 42 | -0.187 | -0.178 | 22:20 | 0.965 |

### Mutation rate

We also investigated potential differences in the mutation rates of island and mainland species. In this study we inferred the mutation rate from dS, the number of synonymous substitutions, along the lineages leading to the mainland and island species (and where there were multiple island and mainland species, from their averages). *N_e_* is predicted to influence mutation rate, and as we found no consistent differences in *N_e_* between island and mainland species we do not expect mutation rate to differ between the two groups. This is in fact the case: comparing dS values between island and mainland species revealed no significant trend (Table 6)(n = 112, p = 0.251). However, when different genomes were considered separately, there was one statistically significant difference between island and mainland species for nuclear DNA (n = 32, p = 0.004). The trend in this instance was for mainland species to have higher values of dS than island species.

**Table 6.** Differences in synonymous divergence (dS) between island and mainland species. The number of comparisons used in each analysis is given in the second column (n), with the significance level of the Wilcoxon signed-ranks test given in the last column. Each particular species comparison appears only once in each dataset.

| Dataset | n | Mean Island dS | Mean Mainland dS | Ratio Island:Mainland | p-value |
| --- | --- | --- | --- | --- | --- |
| Combined | 112 | 0.351 | 1.15 | 52:59 | 0.251 |
| Chloroplast | 10 | 0.0164 | 0.0129 | 6:4 | 0.646 |
| Mitochondrial | 96 | 0.559 | 1.42 | 48:48 | 0.527 |
| Nuclear | 32 | 0.0582 | 0.157 | 6:22 | 0.004 |

## Discussion

It is generally assumed that island species will have smaller effective population sizes than mainland species. Island species are expected to have low effective population sizes initially because they are likely to be founded by a small number of individuals (one pregnant female is sufficient) and hence experience a bottleneck. We find some evidence for this: some island species, which are very closely related to their mainland counterparts, have little or no diversity, consistent with these species experiencing extreme bottlenecks during colonisation. However, besides these species, island species have similar levels of diversity to mainland species. There is no evidence to suggest that island species have low long-term effective populations sizes, despite the fact that island species occupy considerably smaller ranges than mainland species; in this analysis, island species had ranges of on average 0.14% of the area of their mainland counterparts. Consistent with island and mainland species having similar effective population sizes, we find no evidence that natural selection is less efficient in island species.

Our results are perhaps not surprising. It is well established that the relationship between population size and genetic diversity is not straightforward, with levels of genetic diversity remaining remarkably constant across groups of organisms which are incredibly disparate in terms of population size (Leffler et al. 2012; Gillespie & Ohta 1996). What is unique about the current data is that only closely related species are compared to each other-many of the island and mainland species pairs are in the same genus. They therefore share life history traits, many of which influence molecular evolution. In addition, our paired study design allows us to correct for phylogenetic effects (Lanfear et al. 2010). This is crucial, as it has been well demonstrated that molecular evolution is influenced by taxonomy. For example, Romiguier et al. (Romiguier et al. 2014) demonstrated that levels of diversity differ between families but are similar within a family. Correcting for phylogenetic effects has allowed us to study the effects of island colonisation on molecular evolution across a wide range of taxa.

There are a number of possible reasons why island species might not have lower effective population sizes than their mainland counterparts. First, it is possible that many island species are founded by multiple individuals, and gene flow is maintained as they speciate, thereby allowing island species to inherit much of the variation of the mainland species. We have evidence that this is true of some species: birds in particular appear to experience relatively few bottlenecks as a taxonomic group, which is probably a due to their increased dispersal ability relative to other animals. However, after the initial colonisation event, we might then expect the genetic diversity of island species to decline due to their restricted range. We see no evidence of this: even if we exclude those young island species with no diversity, the correlation between synonymous nucleotide diversity and synonymous divergence remains positive (r = 0.214, p = 0.150).

Second, it has been suggested that levels of diversity are relatively constant across species because of an inverse relationship between population size and the mutation rate per generation (Lynch 2007; Piganeau & Eyre-Walker 2009), a relationship for which we have some evidence (Lynch 2010; Sung et al. 2012). This is hypothesised to occur because populations with large effective population sizes can more effectively select for modifiers of the mutation rate. Therefore, selection to reduce the mutation rate will be more effective in larger populations, resulting in lower mutation rates and hence levels of genetic diversity similar to those found in small populations. There is no evidence that this is the case in this analysis. When we analysed the levels of synonymous divergence, an indicator of the neutral mutation rate, we did not find a difference between island and mainland species, indicating that island species do not have higher mutation rates. In addition, there is no evidence, from considering the efficiency of selection, that island species have lower effective population sizes. Finally, upon excluding those species with no diversity we do not find that diversity increases with divergence, which we might expect if higher mutation rates evolve over time in island species.

Third, it is also possible that there is selection on synonymous mutations, which could obscure a relationship between genetic diversity and effective population size. If selection acts on synonymous codons to optimise the accuracy of translation, we expect there to be a distribution of fitness effects of synonymous mutations (Akashi 1994; Stoletzki & Eyre-Walker 2007). We therefore might find that as *N_e_* increases, the proportion of effectively neutral mutations would decrease as selection becomes more efficient. This process could allow the levels of genetic diversity to remain constant as effective population sizes increase, but only if the distribution of fitness effects of synonymous mutations is exponential (Welch et al. 2008). There is no evidence to suggest that this is the case, and therefore this is an unlikely explanation of our results.

Fourth, it has been suggested that levels of genetic diversity might not be correlated to population size due to selection at linked sites (Gillespie 2000; Maynard Smith & Haigh 1974). Gillespie has argued that if the rate of adaptive evolution is mutation limited then as population sizes increase so does the rate of adaptive evolution and hence the level of genetic hitch-hiking – a phenomenon that he has termed genetic draft. Some authors have found evidence to suggest that draft has an important role in reducing genetic diversity. However, studies generally report that draft has relatively weak effects, which may not be powerful enough to reduce genetic diversity to observed levels (Gossmann et al. 2011; Weissman & Barton 2012; Andolfatto 2007). Furthermore, there is no evidence in our data that draft is important. Firstly, if genetic draft was prevalent in our dataset we might expect different patterns for the organellar genomes, which have little or no recombination, and the nuclear genome (Campos et al. 2014). However, they behave qualitatively in a similar fashion. Secondly, we do not find a significant difference between island and mainland species in terms of their DoS. If selective sweeps were responsible for the low diversity of mainland species, we might expect mainland species to have greater values of DoS than their island counterparts. In addition, our results indicate that it is deleterious mutations that are dominating evolutionary dynamics, rather than advantageous mutations. However, it is worth noting that the signal of adaptive evolution could be obscured by a shift in the distribution of fitness effects for island species. Correcting for this with the current dataset is difficult due to a lack of sufficient polymorphism data (we have very few datasets which contain more than 4 alleles), although the results from our limited sample indicate that it is island species that undergo a greater degree of adaptive evolution, rather than species with large population sizes.

Romiguier et al. (Romiguier et al. 2014) recently showed that geographic factors likely to influence population size are poor correlates of genetic diversity when diversity is considered across the full breadth of the animal kingdom. Surprisingly, they find that propagule size is the single best predictor of diversity. Those species with few large propagules had low genetic diversity, and those with a large number of small propagules had high genetic diversity, and were termed K and r strategists respectively. However, the authors do not present a clear hypothesis as to why these strategies should affect genetic diversity. One possibility is that propagule size is related to population density, and that the variance in population density is far greater than the variance in population range size, so that the degree to which species differ in effective and census population sizes is largely determined by density and not range size. However, our results would tend to suggest that population density is not the missing factor, because there is no reason to believe that densities differ systematically between island and mainland species.

Alternatively, it may be that the mutation rate itself is an important determinant of diversity, particularly in organellar genomes (Lynch et al. 2006; Nabholz, Mauffrey, et al. 2008; Bazin et al. 2006). Although the issue is controversial, Nabholz et al. showed that mutation rate is a major determinant of mitochondrial diversity, and as our dataset is dominated by mitochondrial sequences this could explain why we did not find a difference between island and mainland species, considering that we also did not find a difference in mutation rate between them. We found a strong positive correlation between the mutation rate, as measured by the rate of synonymous divergence, and levels of synonymous diversity, both for our entire dataset (n = 138, r = 0.337, p < 0.000), and considering mitochondrial sequences separately (n = 112, r = 0.269, p = 0.004), which lends some support to this theory, however, we are unable to recover this correlation if we correct for phylogenetic independence by comparing island and mainland species (i.e. π_s_(island)/(π_s_(island)+π_s_(mainland)) is not significantly correlated to dS(island) / (dS(island) +dS(mainland)).

In conclusion, our analysis demonstrates that island colonisation typically has little impact on a species’ molecular evolution. For some species the initial colonisation event results in a period of low diversity, but this effect appears to be short-lived with no discernible lasting effects. Our results confirm that census population size is a poor correlate of effective population size.

## Methods

### Dataset

The dataset was created by combining all of the independent island-mainland species comparisons used in two previous studies: the 48 comparisons of island and mainland bird species used in (Wright et al. 2009), and the 44 comparisons used in (Woolfit & Bromham 2005), which cover a wide range of taxa. This dataset was then expanded using a keyword search (‘endemic’) of the Arkive species database (http://www.arkive.org/). One or more mainland relatives and outgroup species were then identified for each island species. This added 56 species comparisons to the dataset. Some comparisons contained a single island and mainland species, while some consisted of multiple island and/or mainland species. All phylogenies were checked for agreement with the literature, and apparent direction of colonisation was noted. In addition, the recorded range area of the species used was calculated from IUCN records (IUCN 2014) using ArcGIS. Protein coding sequences were collected from NCBI (www.ncbi.nlm.nih.gov/genbank/). Sequences were collected if there was an orthologous gene available for each of the island, mainland, and outgroup species in a comparison, or if there were multiple sequences of the same loci available for both the island and the mainland species in a comparison. A note was made of whether the sequences were nuclear, mitochondrial or chloroplast. All alignment files and further details of this analysis are available at: http://dx.doi.org/10.6084/m9.figshare.1296151.

### Statistical tests

This study has a paired design, in that each island species/clade is compared to a closely related mainland species/clade, with each comparison occurring only once in each analysis. If a choice had to be made between comparisons (for example, if statistics from both the mitochondrial and nuclear genomes were available for a single comparison) the statistics that corresponded to the longest sequence alignment were used. This decision should reduce sampling error, because longer sequences are more representative than short sequences. Island and mainland species were compared using Wilcoxon signed-ranks tests. This is a paired, non-parametric test that takes into account the direction of the difference between pairs, and gives greater weight to those pairs that are the most different, making it more powerful than a sign test (Sokal & Rohlf 1995).

### Polymorphism data

Polymorphism data was calculated by aligning sequences of the same loci from the same species using a Geneious translation alignment, which was then analysed using our own scripts. A number of statistics were recorded, including nucleotide diversity and number of polymorphisms. If a comparison included multiple island and/or multiple mainland species, average values of each statistic were taken across the species. Similarly, if multiple sequences from the same genome were available for a particular island/mainland comparison, the average value of the sequences was used. Therefore, each comparison is represented by a single island, mainland, and outgroup value of each polymorphism statistic for a particular genome.

The data was used to calculate π_n_/(π_n_+π_s_), where π_n_ is nonsynonymous diversity and π_s_ is synonymous diversity. This ratio is used because, unlike polymorphism counts, nucleotide diversity is unaffected by the number of chromosomes sampled. In addition, using total diversity as the denominator reduces the number of undefined ratios. Any comparisons with undefined values were excluded from the analysis.

### Substitution data

Substitution data was calculated by aligning orthologs of island, mainland and outgroup species. If multiple sequences at different loci were available for all of the species in a comparison, sequences were concatenated prior to alignment; however, sequences from different genomes of the same organism were treated separately. The alignments were pruned so that they included equal numbers of island and mainland species to control for the node-density effect (Hugall & Lee 2007), and then used to generate phylogenetic trees with RaxMl (Stamatakis 2014), in combination with PartitionFinder (Lanfear et al. 2012). The trees were subsequently used to run the codeml progamme of PAML version 4.7 (Yang 2007), which calculated ω (dN/dS) for island, mainland, and outgroup branches of each tree, as well as separate dN and dS values for each branch.

### Adaptive evolution tests

Polymorphism and substitution data was combined to test for differences in levels of adaptive evolution between island and mainland species. The direction of selection (DoS) statistic was used, calculated as: DoS =dN/(dN+dS) – pN/(pN+pS) This statistic has the advantage over using the neutrality index in that it is defined for all datasets in which there is at least one substitution and one polymorphism, so fewer species comparisons had to be excluded; it is also expected to be unbiased (Stoletzki & Eyre-Walker 2011). Positive values indicate that the dynamics of evolution are dominated by positive selection and negative values that slightly deleterious mutations predominate.

